# The lag phase of seed development plays an important role in determining the maximum potential final seed weight in soybean (*Glycine max* L.)

**DOI:** 10.1101/2025.10.07.679780

**Authors:** Ashwini Shivakumar, Mohammad Foteh Ali, Jacob Sullivan, Montserrat Salmer⍰n, Tomokazu Kawashima

## Abstract

Soybean (*Glycine max* L.) cultivars exhibit substantial variation in seed weight; however, the developmental and physiological mechanisms contributing to this variation remain incompletely characterized. Here, we investigated the relationship between early seed developmental dynamics and final seed weight by comparing two large-seeded and two small-seeded cultivars under control and depodding conditions. Depodding, achieved by retaining a single pod per node, minimized assimilate competition. Final seed weight was positively correlated with cotyledon cell number and duration of the lag phase, which is a key early stage of seed development. Large-seeded cultivars exhibited significantly longer lag phases and higher cotyledon pavement cell numbers than small-seeded cultivars, suggesting that extended lag phases promote enhanced cell proliferation, contributing to increased seed weight. Depodding further increased the cotyledon cell number; however, this response was associated with accelerated embryo development rather than an extension of the lag phase. These findings indicate that both the duration of the lag phase and the rate of early embryo development influence cotyledon cell proliferation and ultimately seed weight. Moreover, genetic factors and assimilate availability regulate developmental processes through distinct pathways. Together, our results highlight the importance of early seed developmental timing in determining final seed weight and provide new insights into the developmental basis of yield-related traits in soybean.

## Introduction

Seed development plays a pivotal role in determining seed yield. The soybean (*Glycine max* L.) seed development process consists of three distinct phases: the lag phase, seed-filling phase, and maturation phase (Fig. 1A). After double fertilization, seed structures such as cotyledons and seed coats are fully formed during the lag phase. At this stage, embryo cells undergo continuous cell division to form preglobular, globular, and heart shapes, and reach the maximum number of cotyledon cells by the end of the lag phase (Egli et al., 1981). The formation of cotyledon cells sets the stage for the subsequent seed-filling phase, where cotyledon cells undergo expansion with the accumulation of reserve materials such as carbohydrates, proteins, and oil, leading to the physical growth of the seeds. The seeds are then desiccated and become quiescent during maturation.

**Figure 1:**
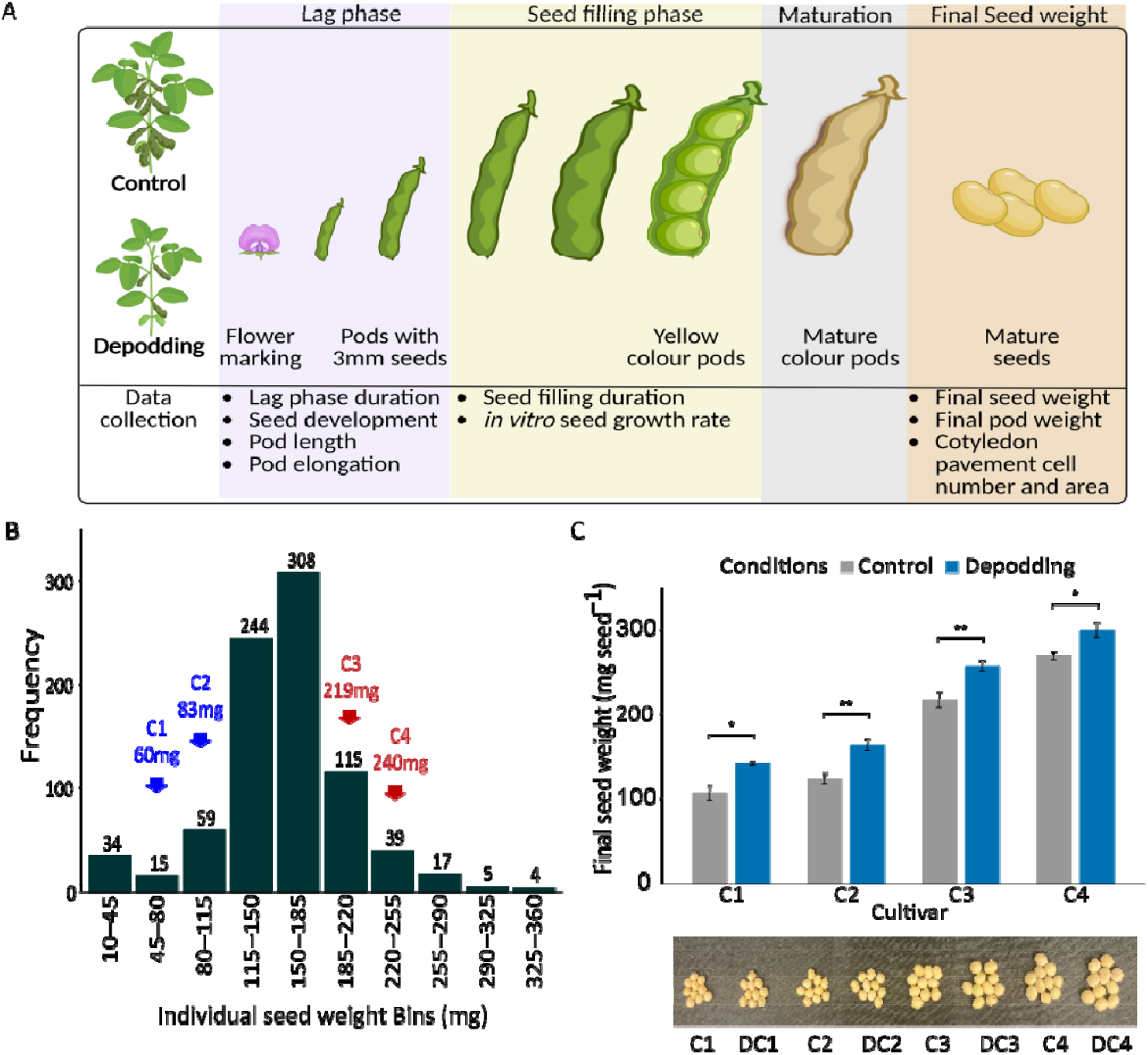
Experimental design and final seed weights of the different cultivars. (A) Experimental design outlining the developmental phases and traits measured under the Control and Depodding conditions. (B) Frequency distribution of seed weight categories in USDA germplasm with MG-0. The X-axis represents the seed weight category bins (mg), and the y-axi represents the frequency distribution of each category. The selected C1-C4 cultivars were selected from the blue- and red-arrowed distribution categories. The seed weights of these cultivars observed by the USDA are mentioned in brackets. (C) Final seed weight (mg) with images of respective seeds below the graph measured for different cultivars (C1, C2, C3, and C4) and conditions (control and depodding). Each experiment and treatment (control and depodding) included five plants per cultivar and was conducted over three consecutive years. Analysis of two-way ANOVA with experiment as a random factor. Tukey’s adjustment with

The weight per seed in crops such as soybean has been gradually optimized over the years of breeding and agronomic practices, resulting in many cultivars with stable and fixed seed weights. However, soybean exhibits significant genetic variation in seed weight, highlighting the potential for further improvements to increase seed weight (Zhang et al., 2024). Despite this variability, it remains unclear how specific stages of seed development contribute to the differences in seed weight. Although seeds remain small (less than 5% of the final weight) during the lag phase, this phase is believed to be important for seed weight because the number of cotyledon cells generated during the lag phase is positively associated with seed weight (Guldan and Brun, 1985a, Egli et al., 1989b, Munier-Jolain et al., 1998). Studies on soybean, maize, and wheat have consistently demonstrated positive correlations between cotyledon/endosperm cell numbers and seed weight, and cotyledon/endosperm cell numbers have also been shown to be positively correlated with the seed growth rate and duration during the seed-filling period (Chojecki et al., 1986, Jones et al., 1996, Guldan and Brun, 1985a). However, other lag phase phenotypes, such as seed development duration and speed, embryo and endosperm development, and their associations with seed weight, remain largely unknown.

The soybean seed weight was strongly influenced by assimilate availability. Reductions in resource supply, such as through shading, have been shown to decrease seed weight, whereas increasing assimilate availability through pod removal leads to increased seed weight (Egli and Bruening, 2001, Chiluwal et al., 2022, Ali et al., 2022). Depodding experiments also showed an increase in cotyledon cell numbers, which enhances seed growth rate and final seed weight (Egli et al., 1989a). However, the broader effects of assimilate supply on seed developmental characteristics and their mechanistic relationships with seed weight remain poorly understood.

Pod development itself is another key factor influencing seed growth. In addition to encapsulating and protecting developing seeds, pods support seed development by supplying assimilates (Bennett et al., 2011). Pod length has been positively correlated with seed size and serves as a reliable predictor of seed weight (Frank and Fehr, 1981, Fraser et al., 1982). Conversely, pods may also constrain seed growth, potentially limiting final seed weight (Egli et al., 1987). Despite these insights, limited information is available on specific pod developmental traits such as pod elongation rate and pod weight, and their direct relationships with seed developmental processes remain largely unresolved.

Thus, our project aimed to address the lack of knowledge regarding seed development phenotypes and assimilate supply in relation to soybean weight at a physiological level as well as any potential interactions between genotypes and responses to assimilate supply. We hypothesized that differences in seed weight might be impacted by the seed developmental period, particularly by the lag phase phenotypes. To test our hypothesis, we conducted experiments using four soybean cultivars, with or without depodding (pod removal). We quantified seed developmental phenotypes, such as embryo development, seed developmental duration (lag phase and seed-filling duration), pod length, pod elongation rate, seed growth rate, cotyledon cell area, cotyledon cell number, and seed weight.

## Results

## Influence of Genotypic Variation and Assimilate Competition on Seed and Pod Development

A notable diversity in seed weight was observed among cultivars in the USDA SOY germplasm maturity group-0 repository (Fig. 1B). To gain insight into the stages, duration, and physiological aspects of seed development by focusing on cultivars with different seed weights, we selected four cultivars from the same maturity group that exhibited differences in seed weight. We focused on small-seeded cultivars, denoted as C1 (PI 607835) and C2 (PI 593655), and their counterparts with larger seeds, namely, C3 (PI 603322) and C4 (PI 594245).

The maximum attainable weight of soybean seeds is constrained by intense competition for photoassimilates among sinks, such as the reproductive organs (Borrás et al., 2004). A strategic approach to mitigating this competition has been observed in previous studies, where depodding (removal of all but one pod per node) was employed as physiological tool to remove assimilate competition and enhance assimilate concentration (Egli and Bruening, 2001, Ali et al., 2022). To ascertain whether assimilate competition contributed to seed weight differences in the selected large- and small-seeded cultivars, we implemented the depodding condition for each individual cultivar. The results revealed a significant increase in the final seed weight of all cultivars in the depodding condition compared to the control (Fig. 1C). This suggests that assimilate supply influences seed weight, making it essential to control for this parameter by depodding when conducting seed size or weight analyses. Despite the introduction of the depodding condition, the cultivar characterized by smaller seeds (C1 - control and depodding) persisted as the smallest, while the cultivar with larger seeds (C4 - control and depodding) remained the largest. These findings indicate that although assimilate supply and competition can modify seed weight, the differences in seed weight among these four cultivars are genetically controlled.

Pod growth and development play important roles in the physical protection of seeds, distribution of the assimilate supply, and physical restriction to seeds in attaining potential seed weight (Crafts-Brandner and Egli, 1987, Fukuta et al., 2005, Egli et al., 1987). To understand how pod physiology could affect differences in seed weight, we monitored pod length and pod elongation rate during the pod developmental stage and obtained the final pod weight at harvest. The pod elongation rate and pod length were significantly higher in the large-seeded cultivars than in the small-seeded cultivars under both control and depodding conditions (Table 1, Supplementary Fig. S1, A and B). No significant differences in pod elongation rates or pod length were observed between the control and depodding conditions. Our results indicate that pod elongation rate and pod length are genetically controlled traits that do not change with the manipulation of the assimilate supply. However, pod weight was significantly increased in most cultivars (C2, C3, and C4) upon depodding (Supplementary Fig. S1C). The increase in pod weight under depodding conditions shows that assimilates may also be loaded to the pod along with seeds, further validating that depodding conditions enhance the assimilate supply and remove competition.

**Table 1:** ANOVA table for the analysis of seed and pod characteristics for cultivars with different seed weights.

| Seed phenotypes | Variables | Cultivar (E) | Treatment (T) | Interaction (E*T) |
| --- | --- | --- | --- | --- |
| Final seed weight | DF | 3 | 1 | 3 |
|  | F Value | 238.2137 | 53.3907 | 0.2218 |
|  | P > F | < 0.001*** | < 0.001*** | 0.88 |
| Lag phase duration | DF | 3 | 1 | 3 |
|  | F Value | 29.1351 | 6.421 | 0.3834 |
|  | P > F | < 0.001*** | 0.02* | 0.76 |
| Seed filling duration | DF | 3 | 1 | 3 |
|  | F Value | 31.268 | 60.179 | 1.5 |
|  | P > F | < 0.001*** | < 0.001*** | 0.27 |
| *In vitro* seed growth rate<br>(mg day<sup>-1</sup>) | DF | 3 | 1 | 3 |
|  | F Value | 20.7083 | 2.9358 | 3.2325 |
|  | P > F | < 0.001*** | 0.08. | 0.02* |
| Cotyledon pavement cell number | DF | 3 | 1 | 3 |
|  | F Value | 391.8343 | 166.8628 | 9.6036 |
|  | P > F | < 0.001*** | < 0.001*** | < 0.001*** |
| Cotyledon pavement cell area | DF | 3 | 1 | 3 |
|  | F Value | 43.8748 | 23.0836 | 2.9575 |
|  | P > F | 2.2E-16*** | 4.426E-06*** | 0.035508* |
| Pod elongation rate | DF | 3 | 1 | 3 |
|  | F Value | 20.4787 | 2.6366 | 3.2104 |
|  | P > F | < 0.001*** | 0.10976 | 0.02* |
| Pod length | DF | 3 | 1 | 3 |
|  | F Value | 72.3671 | 0.8133 | 1.1886 |
|  | P > F | < 0.001*** | 0.3707 | 0.3216 |
| Final Pod weight | DF | 3 | 1 | 3 |
|  | F Value | 70.3329 | 88.9264 | 6.4274 |
|  | P > F | < 0.001*** | < 0.001*** | < 0.001*** |

Significant codes indicated in the table: \*\*\* 0.001; \*\* 0.01; \* 0.05; . 0.1

### The duration of the lag phase is positively associated with seed weight

The duration of the seed development phase is crucial for seed weight and yield (Egli, 2017). Large-seeded cultivars exhibited longer lag phase durations than small-seeded cultivars under both control and depodding conditions (Table1, Fig. 2A). The lag phase duration did not show a significant difference between the control and depodding conditions for all cultivars. Furthermore, the correlation analysis between lag phase duration and seed weight revealed a significant positive relationship under both the control and depodding conditions (Fig. 2C).

**Figure 2:**
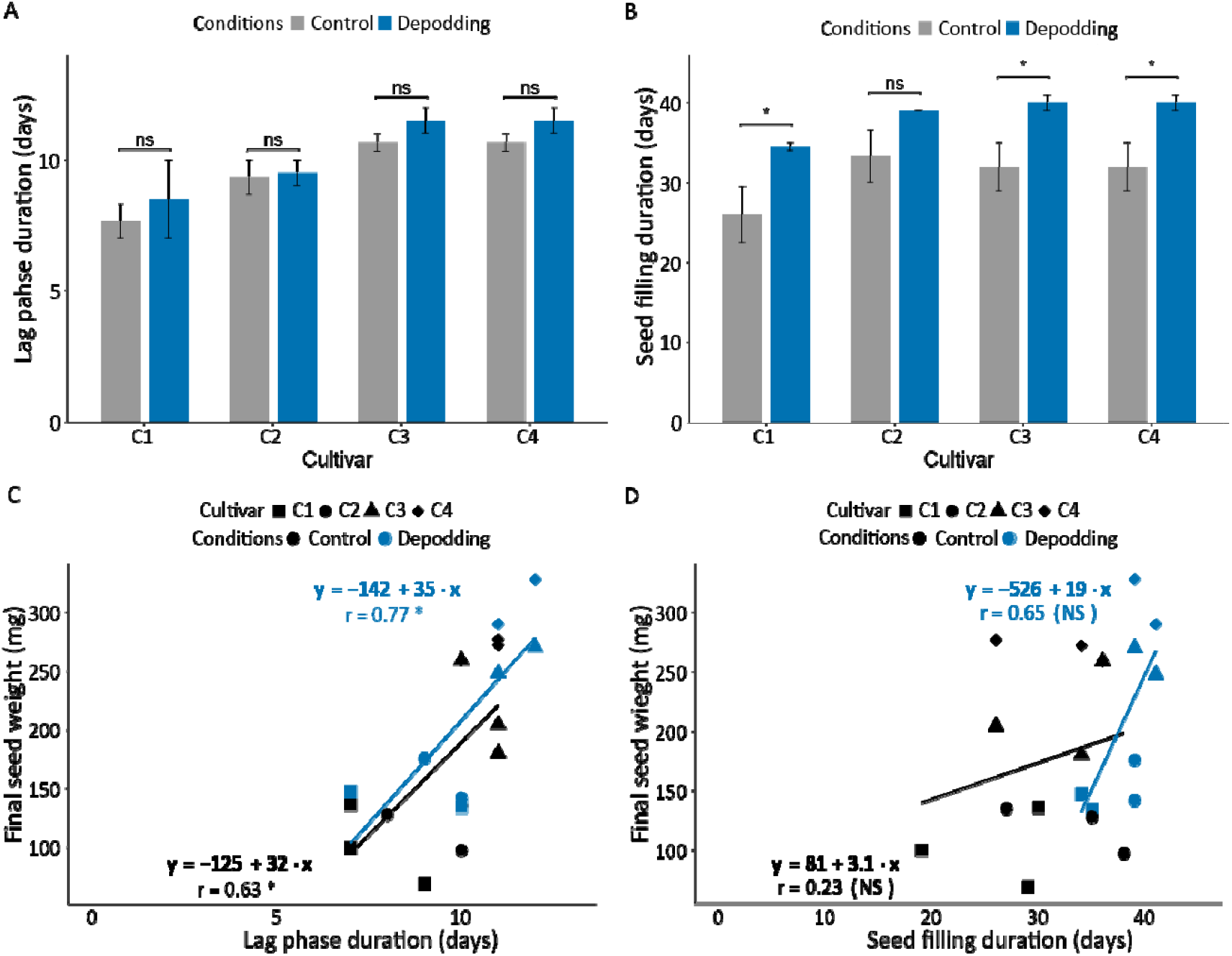
Seed developmental duration of cultivars with different seed weights. (A) Lag phase duration and (B) seed-filling duration (days) measured under the control and depodding conditions. Each experiment and treatment (control and depodding) included five plants per cultivar and was conducted over three consecutive years. Analysis of two-way ANOVA with experiments as a random factor. Error bars represent standard errors. Tukey’s adjustment with contrast was used to determine statistical significance. Linear regression of (C) lag phase and (D) seed-filling duration with seed weight. Black and blue represent control and depodding conditions, respectively. The symbols represent each cultivar (C1-C4). Each data point corresponds to the duration (lag and seed-filling phases) and average seed weight, respectively. The parameters of the fitted linear regression (line) and the correlation coefficient (r) are also shown. (* p ≤ 0.05, ** p ≤ 0.01, *** p ≤ 0.001 indicate the significance; ns-nonsignificant)

We did not observe a significant difference in seed-filling duration between the large- and small-seeded cultivars (Fig. 2B). However, the seed-filling duration showed a significant increase in the depodding conditions compared with the control (Table 1, Fig. 2B). Regression analysis between seed-filling duration and seed weight showed a positive correlation for both the control and depodding conditions; however, the correlation was not significant (Fig. 2D). These results suggest that the lag phase duration is primarily governed by genetic factors and is strongly and positively associated with seed weight. Conversely, seed-filling duration appears to be more dynamically responsive to changes in the assimilate supply.

### Embryo development accelerates in the depodding condition

Given the significant correlation observed between the lag phase duration and seed weight (Fig. 2, A and C), we conducted a detailed investigation of embryo and endosperm development during this phase. The embryo and endosperm in young seeds from 1-2 cm pods during the lag phase from all four cultivars were categorized into five developmental stages: globular, early heart, mid-heart, late-heart, and cotyledon embryos stages (Fig. 3A). No clear differences in embryo growth were observed between cultivars with small and large seeds under either condition (Fig. 3B). However, developmental progression was accelerated under depodding conditions compared with the control across all cultivars. For example, in cultivar C1, 35% of seeds under depodding had reached the late heart embryo stage and 20% had progressed to cotyledonary stage. In the control on the other hand, the majority (42.9%) remained at the mid heart embryo stage, with no seeds advancing to later stages. A similar trend was observed in the other cultivars. These results show that alterations in the assimilate supply contribute to the acceleration of embryo development during the lag phase.

**Figure 3:**
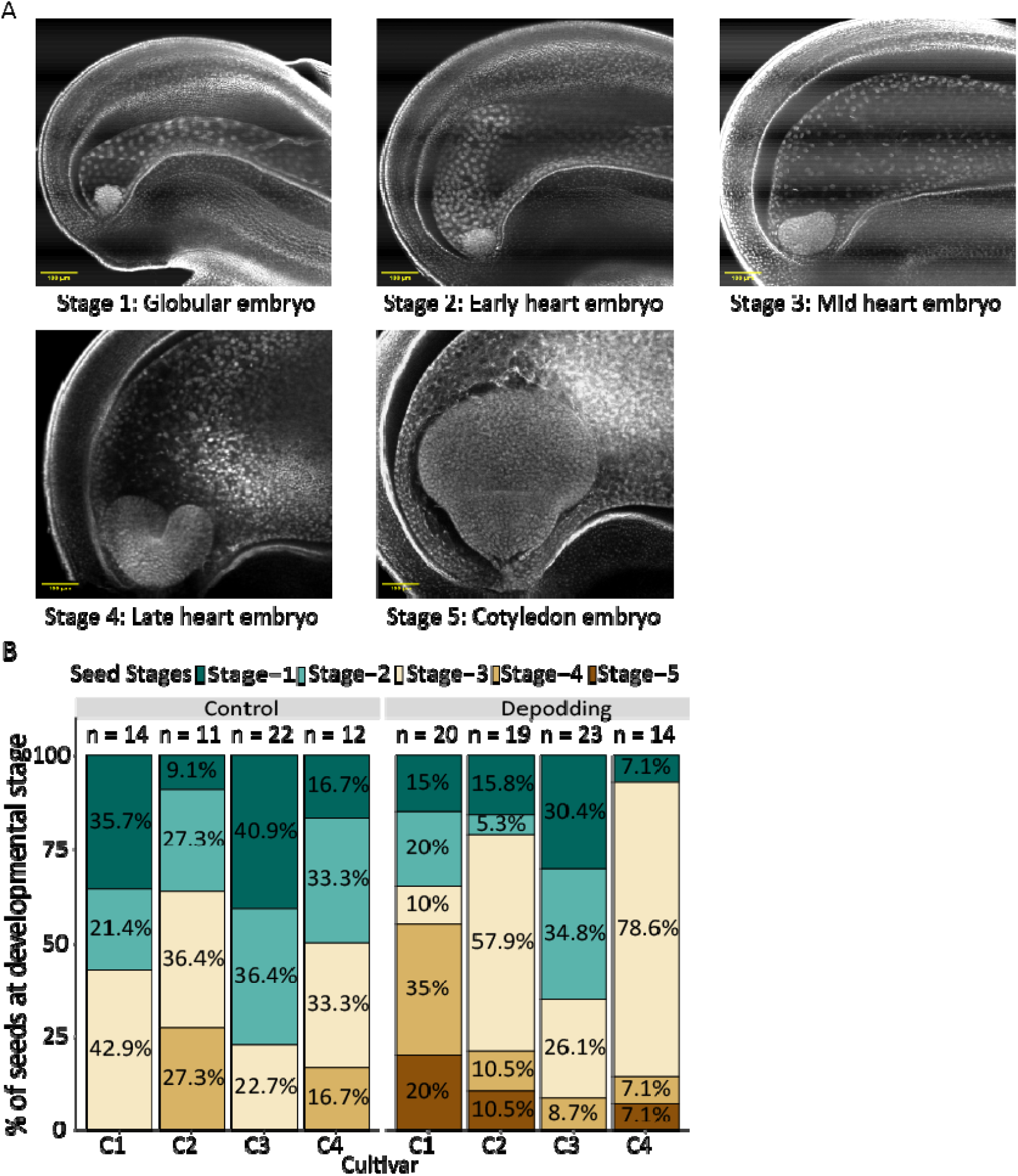
Embryo and endosperm development of cultivars with different seed weights. (A) Confocal images of six stages of embryo development. (B) Embryo development and endosperm of soybean seeds collected from to 1-2 cm pods were observed in experiments 1 and 2 under control and depodding conditions, respectively. The X-axis denotes the cultivars, Y-axis denotes the percentage of different stages of seeds observed, and n denotes the number of seeds used for analysis.

### Seed weight reflects a complex balance between cotyledon cell number and size

The *in vitro* seed growth rate (SGR) or *in vitro* potential seed growth assesses the potential seed growth under conditions with abundant sucrose and amino acids, effectively removing carbon and nitrogen limitations (Egli and Wardlaw, 1980b, Egli et al., 1989a). This method eliminates constraints from pod competition and pod wall restrictions during seed-filling, providing insight into the intrinsic maximum potential for seed growth (Fig. 4A). We observed significant differences in *in vitro* SGR between the small- and large-seeded cultivars under both control and depodding conditions (Table 1, Fig. 4B). Exception for cultivar C2, depodding tended to increase the *in vitro* SGR relative to the control.

**Figure 4:**
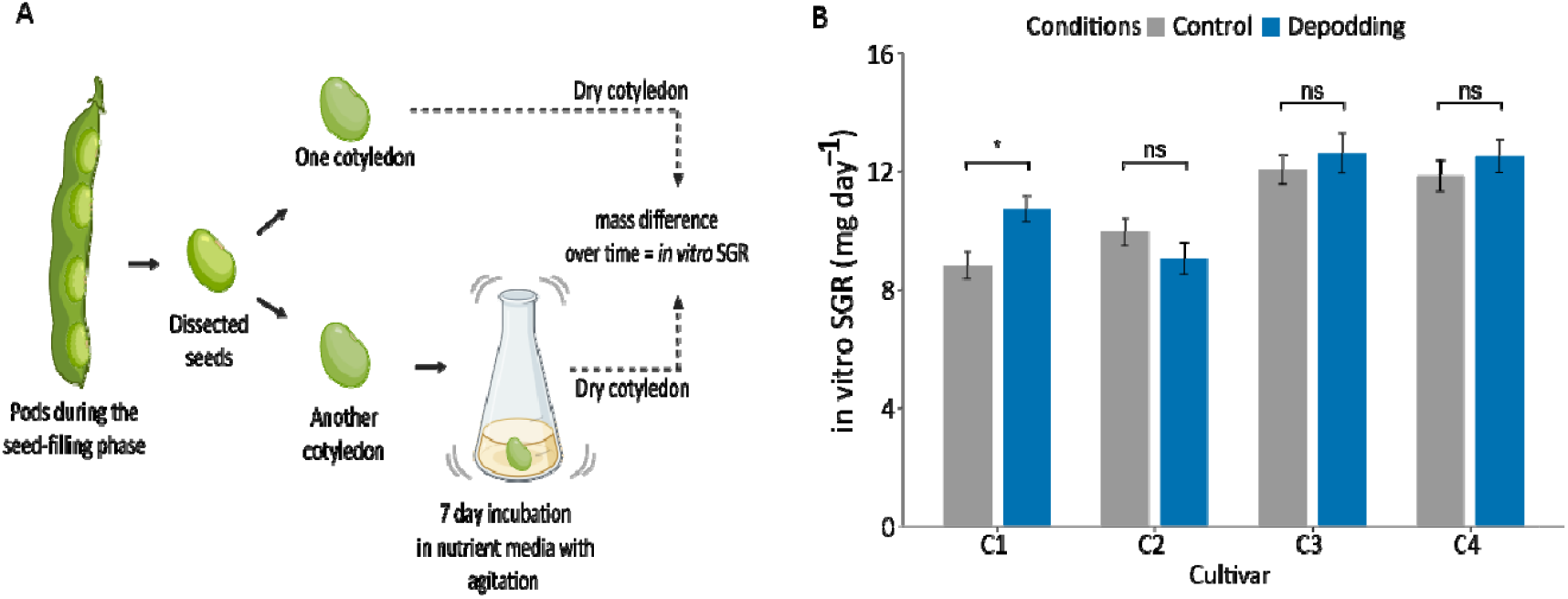
Experimental design and analysis for in vitro seed growth rate. (A) Experimental design outlining the *in vitro* seed growth rate protocol and (B) in vitro seed growth rate (mg day^-1^) of different cultivars (C1, C2, C3, and C4) and conditions (Control and Depodding). Each experiment and treatment (control and depodding) included five plants per cultivar and was conducted over three consecutive years. Analysis of two-way ANOVA with experiments as a random factor. Tukey’s adjustment with contrast was used to determine statistical significance. (* p ≤ 0.05, ** p ≤ 0.01, *** p ≤ 0.001 indicate the significance; ns-nonsignificant)

Cotyledon cell number, established during the lag phase, was positively correlated with both *in vitro* SGR and final seed weight (Egli et al., 1989a, Guldan and Brun, 1985b). To further investigate this relationship, scanning electron microscopy was used to quantify pavement cell number and area on the ventral surface of the mature soybean cotyledons (Fig. 5C). We observed a significant increase in cotyledon cell numbers in the large-seeded cultivars opposed to small-seeded cultivars, as well as in the depodding condition compared to control (Table 1, Fig. 5A). These findings imply that an increase in cotyledon cell number contributes to a greater seed weight. However, cotyledon cell area showed an inverse pattern (Fig. 5B), decreasing as cell number increased although the overall seed weight aligned more closely with cell number. This apparent trade-off highlights the complexity of seed weight determination, which depends not only on the total number of cotyledon cells but also their individual growth dynamics. Consequently, final seed weight results from a delicate balance between cell proliferation and expansion, rather than a simple additive effect of cell number alone.

**Figure 5:**
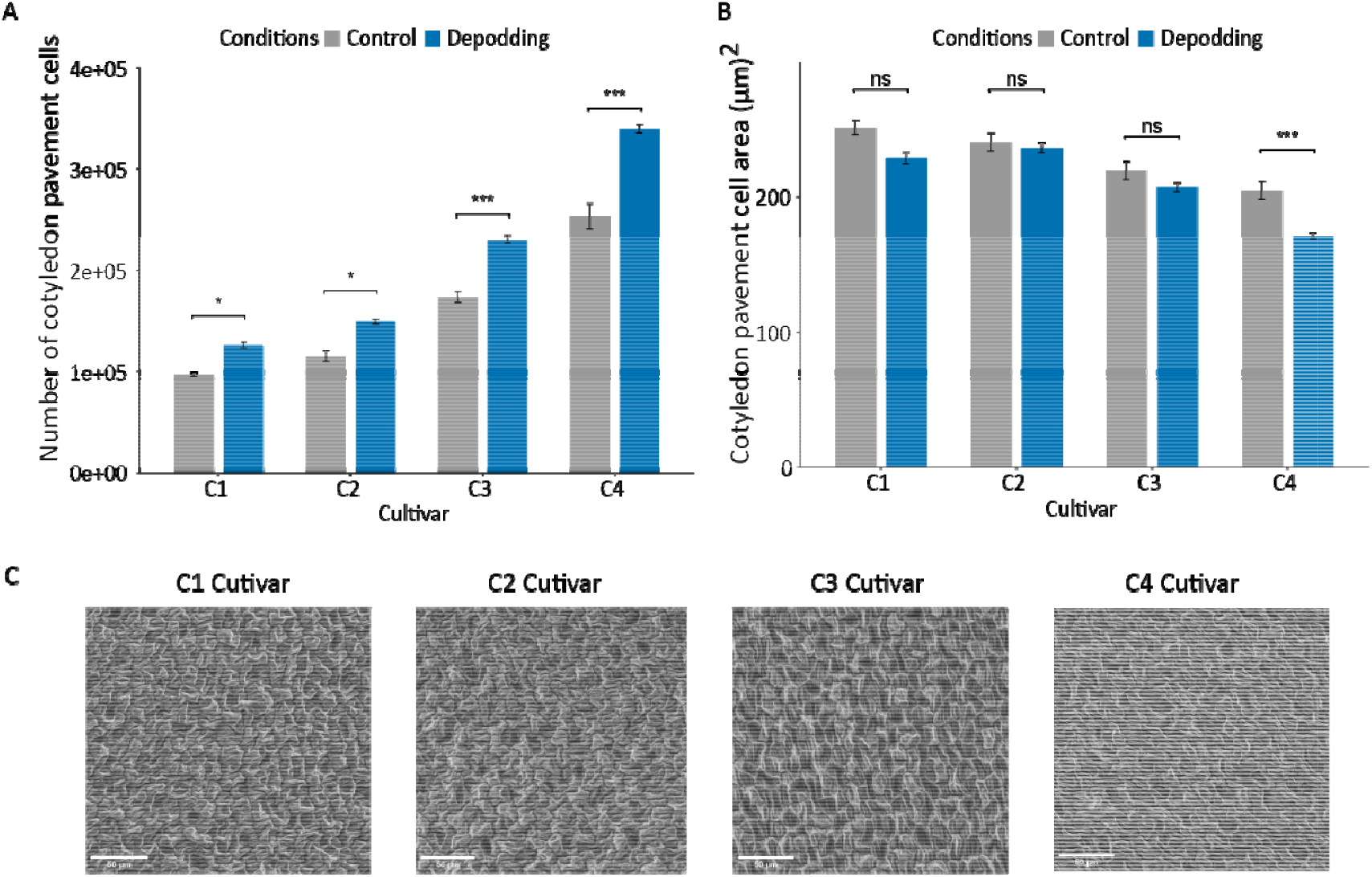
Cotyledon pavement cell numbers and cell area of different seed weight cultivars. (A) cotyledon pavement cell number, (B) cotyledon pavement cell area, and (C) scanning electron microscope images of the ventral surface of the cotyledons of different cultivars (C1, C2, C3, C4) and conditions (Control and Depodding). Each experiment and treatment (control and depodding) included five plants per cultivar and was conducted over three consecutive years. Analysis of one-way ANOVA and Tukey’s adjustment with contrast was used to determine statistical significance. (* p ≤ 0.05, ** p ≤ 0.01, *** p ≤ 0.001 indicate the significance; ns-nonsignificant)

## Discussion

This study provides new insights into the physiological and developmental mechanisms underlying seed weight variation in soybean. By comparing large- and small-seeded cultivars under both control and depodding conditions, we identified key processes contributing to final seed weight. Our findings highlight two distinct regulatory pathways clearly: one governed by assimilate availability and the other by genetic programming (Fig. 6). Assimilate supply primarily modulates the rate and duration of resource-driven growth, accelerating processes such as early embryogenesis and extending seed filling in a genotype-independent manner. By contrast, genetic programming governs the intrinsic timing of developmental phases, particularly the lag phase, which sets the baseline window for cell proliferation and thereby establishes a cultivar’s inherent seed weight potential. Although these pathways converge on cotyledon cell proliferation as a common outcome, their mechanisms are clearly separable: one external and environmentally responsive, the other intrinsic and developmentally programmed. Previous studies often treated these influences as intertwined, but our analysis reveals that they operate through independent biological routes.

**Figure 6:**
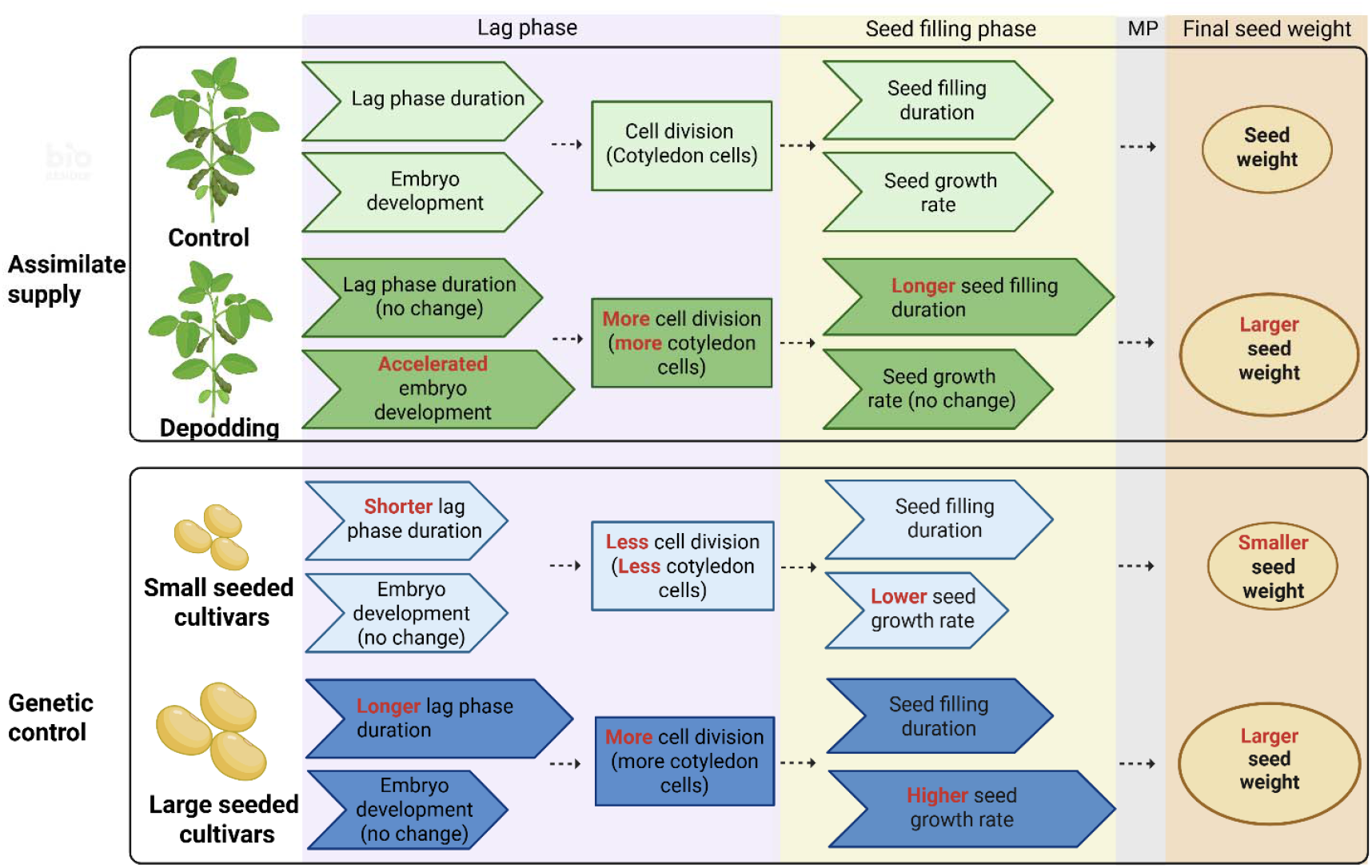
Assimilate supply and genetic pathways controlling seed size. The schematic diagram illustrates the differences in seed development under varying assimilate supply conditions (control and depodding) and genetic backgrounds (small- and large-seeded cultivars). In the assimilate supply pathway, the control condition (top row) represents embryo and seed development characteristics under standard conditions, serving as a reference. By contrast, depodding (second row) accelerates embryo development, leading to increased cotyledon cell number, extended seed-filling duration, and greater seed weight compared to control. Genetically, the small-seeded cultivar (third row) exhibits a shorter lag phase, producing fewer cotyledon cells, lower in vitro seed growth rate (SGR), and reduced final seed weight. The large-seeded cultivar (bottom row), however, displays a prolonged lag phase, enabling greater cell proliferation, higher *in vitro* SGR, and increased seed weight. The lag and seed-filling phase are indicated at the top of the diagram to represent the developmental timeline across the conditions.

In the assimilate supply pathway, increased resource availability accelerated embryo development (Fig. 3B) likely by promoting early cell proliferation and increasing cotyledon cell number (Fig 5A). During rapid cell division, embryos require a high sugar supply, not only as nutrient source providing carbon and energy, but also as a signaling molecule regulating cell cycle progression (Wobus and Weber, 1999, Ruan et al., 2010, Ruan et al., 2012). Sugar availability has been shown to stimulate the expression of cyclin genes (e.g., *CycD2* and *CycD3*), key regulators of the G1 phase (Riou-Khamlichi et al., 2000, Wang and Ruan, 2012, Wang and Ruan, 2013). Consistently, high sugar levels correlate with increased cyclin gene expression in *Vicia faba* embryo (Weber et al., 1996) and Arabidopsis endosperm (Wang and Ruan, 2012). Together, in our depodding conditions, the higher assimilate supply during embryo development may therefore accelerate cell proliferation by providing both metabolic energy and coordinated regulation of cell cycle genes. Depodding also extended the seed-filling period (Fig. 2B), further contributing to increased seed weight. Notably, the seed-filling duration can be extended by depodding at different times during this phase (Chiluwal et al., 2022), suggesting that its duration is not strictly determined during the lag phase or early seed development. Instead, it is likely a consequence of increased assimilate availability at the time of depodding, an outcome rather than a causal factor in seed size. Further studies are needed to clarify how seed-filling duration is regulated. Importantly, both small- and large-seeded cultivars responded similarly to enhanced assimilate supply, indicating a consistent, genotype-independent effect. This pathway primarily influences seed development by modulating the rate and duration of resource-driven growth, particularly during early embryogenesis.

By contrast, the genetically regulated pathway shapes seed development through intrinsic developmental timing. Large-seeded cultivars exhibited a consistently longer lag phase than small-seeded ones (Fig. 2A), providing an extended window for cell proliferation during early embryogenesis. This led to higher cotyledon cell numbers and was strongly correlated with both final seed weight (Fig. 2C) and *in vitro* SGR (potential seed growth). Similar positive correlation of seed weight with cotyledon cell numbers was observed in *Pisum sativum* (Lemontey et al., 2000) and *Brassica napus* (Li et al., 2015). However, under depodding conditions, although cotyledon cell number increased, *in vitro* SGR remained unchanged (Fig. 4B), suggesting that assimilate-driven proliferation does not always translate into enhanced intrinsic growth potential. Interestingly, cotyledon cell size exhibited an inverse trend relative to cell number (Fig. 5A and 5B). While large-seeded cultivars had more cells, these cells were generally smaller, indicating a compensatory mechanism that may preserve overall cotyledon volume. This trade-off points out the complexity of seed weight determination, where both cell proliferation and expansion are finely coordinated. Final seed weight is not simply the result of increased cell number; rather, it reflects the interplay between the extent of cell division and the growth capacity of individual cells.

Taken together, our study disentangles two fundamentally distinct regulatory pathways that shape seed development and final seed weight. By dissecting these pathways, we not only clarify the mechanistic basis of seed weight variation in soybean but also provide a framework for future breeding strategies that strategically target either assimilate responsiveness or intrinsic developmental timing to enhance yield potential.

## Materials and methods

### Experimental design

Three greenhouse experiments were conducted at the University of Kentucky, Lexington, Kentucky (38.02° N, 84.50° W) in 2019 (May 20^th^), 2021 (June 24^th^) and 2022 (June 28^th^). Four soybean cultivars of maturity group 0 (MG-0) were selected for different seed sizes from the SOY repository of the US National Plant Germplasm System (Fig. 1B). Seeds were inoculated with *Bradyrhizobium japonicum* (Advanced Biological Marketing, Van Wert, OH) and sown in 3.8 L pots (one plant per pot) filled with a 6:1 ratio of pro-mix growing medium and sterilized loam soil. Twelve grams of slow-release fertilizer (Osmocote, 14:14:14, N:P: K) were provided to each pot.

The experimental design consisted of a completely randomized design with source-sink manipulation conditions, namely control and depodding. Each experiment and treatment (control and depodding) included five plants per cultivar and was conducted over three consecutive years. Open flowers in the control and depodding conditions on the main stem were marked with acrylic paint to maintain uniformity in flower/pod age. Simultaneously, depodding was performed at an open flower stage on the main stem by removing all flowers, except one marked flower per node. Approximately all the flowers on the secondary branches were removed during depodding. All flowers that developed after depodding were continuously removed throughout the experiment.

### Data collection

Marked flowers/pods from each experiment were used for phenotypic observations. The pod developmental stages were recorded every three days until maturity. The duration of the lag phase was defined as the number of days from open flower to pods with 3 mm seeds, and the duration of the seed-filling phase was defined as the number of days from pods with 3 mm seeds to yellow pods (Nico et al., 2015, Ali et al., 2022). Pod length was recorded on alternate days with electronic calipers from 5 mm pods to the end of the lag phase (pods with 3 mm seeds). At full plant maturity, R8 (Fehr and Caviness, 1977) marked pods were harvested separately, counted, and dried at 65 °C for 48 h. The pod weight, number of seeds, and final seed weight were collected from the marked pods and the individual seed weights were calculated.

### Microscopic assays

For the microscopic assay, seeds were collected from marked pods (1-2 cm pod length) in 2019 and 2021. The pods were collected on the same day and cut open to remove seeds, which were then placed in a fixing solution (EtOH/acetic acid, 3:1) and stored at 4 °C until processing. The seeds were processed using the Feulgen staining method (Braselton et al., 1996), with propidium iodide (PI) added as a modification of this procedure. After mounting in LR resin, embryo and endosperm development was observed using a confocal microscope with settings following those used by (Ali et al., 2022). All confocal images were analyzed and processed using the Fiji (ImageJ) software.

### *in vitro* seed growth rate assays

To examine the *in vitro* seed growth rate (SGR), marked pods were collected from four to five plants during the seed filling phase and cultured (Egli and Wardlaw, 1980a). After washing the harvested pods with a mixture of liquinox soap and water, the seeds were carefully extracted from the clean pods under sterile conditions. The seed coat was removed, and the embryonic axis was carefully separated. One cotyledon from each seed was placed in a 50-ml Erlenmeyer flask with 7 ml of culture solution (Egli and Wardlaw, 1980a, Ali et al., 2022) and incubated for 7 days with shaking at room temperature. The other cotyledon from the same seed was dried to determine the initial dry weight. After incubation, the cotyledon was removed from the flask and dried to determine the final dry weight. Contaminated flasks were discarded and the *in vitro* seed growth rate (mg seed^-1^ d^-1^) was calculated as the difference between the initial and final dry weights of each pair of cotyledons.

### Cotyledon cell numbers

Seeds harvested (at R8 stage) from the four cultivars and two conditions (control and depodding) from 2022 were used to excise cotyledons. The ventral surfaces of the cotyledons were examined to observe the cotyledon pavement cells. Observations were conducted using a scanning electron microscope (FEI Quanta 250) under low-vacuum conditions with water (Fig. 4C). The imaging was performed at the Electron Microscopy Center of the University of Kentucky. Following imaging, the cotyledon cell size and counts were analyzed using the ImageJ software.

### Statistics

Data from lag phase duration, seed filling duration, *in vitro* SGR, pod elongation rate, pod length, individual pod weight, and individual seed weight were analyzed by condition (control and depodding) and their interaction as fixed effects and year as random effects using “lmertest” package in RStudio4.1. Cotyledon pavement cell number from 2022 was analyzed by ANOVA with conditions (control and depodding) and their interaction as fixed effects using “lm” function R package. Tukey’s correction with contrasts was used to generate significance between the conditions (* *p* ≤ 0.05, ** *p* ≤ 0.01, *** *p* ≤ 0.001) (Supplementary Table S2). Graphs were developed using the R package ggplot2.

## Acknowledgements and Funding

We thank Dr. Egli for reading the manuscript and for providing comments and suggestions. We thank Dr. Nicolas Briot and Dr. Deepak Kumar for helping to acquire the scanning election microscope (SEM) images and the Electron Microscopy Center, College of Engineering, University of Kentucky for providing the SEM. Artificial intelligence-based language correction software was used throughout the text to ensure accurate spelling and grammar as well as to achieve the most comprehensible style possible without influencing the content. Images were generated using BioRender (https://BioRender.com) and Adobe Illustrator.

Ashwini Shivakumar was supported by a Ph.D funding Netaji Subhas ICAR (Indian Council of Agriculture Research) international fellowship, India. Tomokazu Kawashima was supported by the Division of Molecular and Cellular Biosciences, award no. 2334516, from the National Science Foundation, and by the W5168 Hatch Multistate research capacity funding program from the USDA National Institute of Food and Agriculture.

## Author Contributions

A.S., M.F.A., M.S. and T.K. conceived and designed the study. A.S., M.F.A., and J.S. performed experiments. A.S. and M.F.A. analyzed the data. A.S., M.S., and T.K. interpreted data. A.S. and T.K. wrote the paper. All the authors have read and approved the final manuscript.

---

**Supplementary Table S1:** Contrast analysis for seed and pod characteristics for different seed wight cultivars. Final seed weight (FSW), pod elongation rate (PER), pod length (PL), final pod weight (FPW), lag phase duration (LP), seed-filling duration (SFD), in vitro seed growth rate (SGR), and cotyledon pavement cell number (CPN).

| Variables | FSW | PER | PL | FPW | LP | SFP | SGR | CPN |
| --- | --- | --- | --- | --- | --- | --- | --- | --- |
| C1 - C2 | 0.0406 | 0.68 | 0.9999 | 0.0003 | 0.0247 | 0.0001 | 0.878 | 0.0059 |
| C1 - C3 | <.0001 | <.0001 | <.0001 | <.0001 | 0.0001 | 0.0001 | <.0001 | <.0001 |
| C1 - C4 | <.0001 | <.0001 | <.0001 | <.0001 | 0.0001 | 0.0001 | <.0001 | <.0001 |
| C2 - C3 | <.0001 | <.0001 | <.0001 | <.0001 | 0.0063 | 0.9956 | <.0001 | <.0001 |
| C2 - C4 | <.0001 | 0.0001 | <.0001 | <.0001 | 0.0063 | 0.9956 | <.0001 | <.0001 |
| C3 - C4 | <.0001 | 0.9989 | 0.7309 | 0.9968 | 1 | 1 | 0.9904 | <.0001 |

**Supplementary Figure S1:**
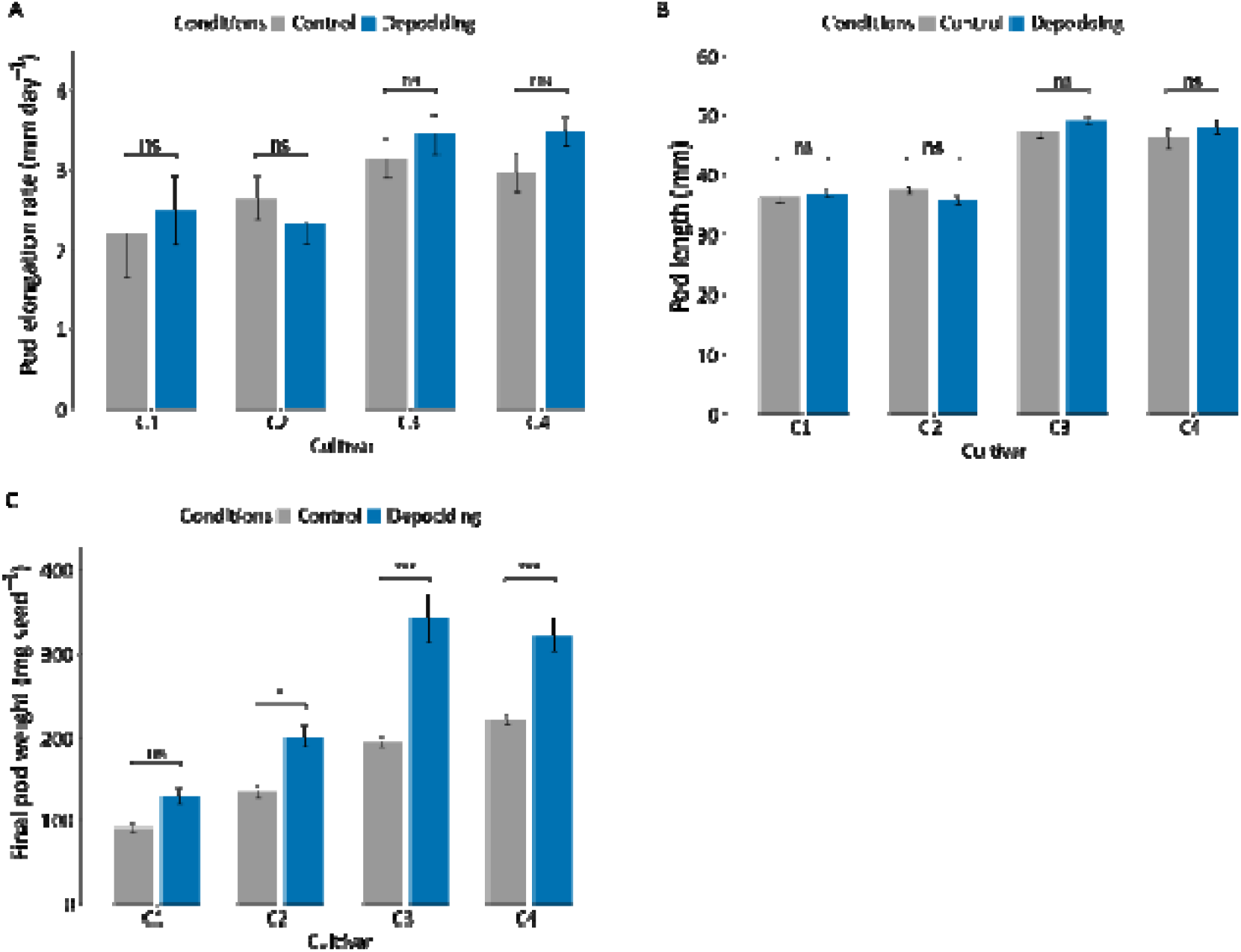
Pod characteristics of different seed weight cultivars. (A) Pod elongation rate (cm day^-1^), (B) Pod length (mm), (C) final pod weight (mg) measured for different cultivars (C1, C2, C3, C4) and conditions (control and depodding). Analysis of two-wa ANOVA with experiments (Experiment 1-3) as a random factor. The tukey adjustment with contrasts was used to determine statistical significance. (* p⍰≤⍰0.05, ** p⍰≤⍰0.01, *** p⍰≤⍰0.001 indicate significance; ns-nonsignificant).

